# Optimal construction of army ant living bridges

**DOI:** 10.1101/116780

**Authors:** Jason M. Graham, Albert B. Kao, Dylana A. Wilhelm, Simon Garnier

## Abstract

Integrating the costs and benefits of collective behaviors is a fundamental challenge to understanding the evolution of group living. These costs and benefits can rarely be quantified simultaneously due to the complexity of the interactions within the group, or even compared to each other because of the absence of common metrics between them. The construction of ‘living bridges’ by New World army ants - which they use to shorten their foraging trails - is a unique example of a collective behavior where costs and benefits have been experimentally measured and related to each other. As a result, it is possible to make quantitative predictions about when and how the behavior will be observed. In this paper, we extend a previous mathematical model of these costs and benefits into a general framework for analyzing the optimal formation, and final configuration, of army ant living bridges. We provide experimentally testable predictions of the final bridge position, as well as the optimal formation process for certain cases, for a wide range of scenarios, which more closely resemble common terrain obstacles that ants encounter in nature. As such, our framework offers a rare benchmark for determining the evolutionary pressures governing the evolution of a naturally occurring collective animal behavior.

## 1. Introduction and Background

Over the past four decades, studies have revealed the functional consequences of collective animal behaviors, which are often driven by interactions between individuals with little or no global knowledge [1, 2, 3, 4, 5]. The cohesive movement of bird flocks and fish schools, some of the most visually striking examples of how animal groups can dynamically self-organize, allow for improved migration accuracy [6], predator avoidance [7], and resource finding [8]. However, collective behavior also operates at less conspicuous, but equally functionally important, scales, in order to generate division of labor [9], pattern formation [10, 11], or physical construction [12, 13] across many animal taxa.

One of the principal challenges in studying collective behavior is simultaneously quantifying both the benefits and costs associated with group living in order to understand the overall selective pressure on the behavior and hence its evolution. In some cases, the benefits, such as the improvements in navigation [14], or the costs, such as an increase in the risk of disease [15], have been measured in isolation. However, since the proximate currencies of fitness related to benefits and costs are often very different (e.g. navigation direction and disease risk), and operate at different spatial or temporal scales, it is often difficult to measure both benefits and costs simultaneously in order to estimate the ultimate fitness consequences of group living.

The construction of living bridges by the army ant *Eciton hamatum* is a unique example of a collective behavior that is amenable to measurements of both costs and benefits [16]. Therefore, it allows for quantitative predictions about when and how the behavior will be observed. Found in the tropical forests of Central and South America, army ants are nomadic, moving their entire colony (sometimes exceeding a million individuals) to a new location each day in search of new sources of food while the colony has developing young [17, 18, 19]. As a consequence of this nomadic lifestyle, these ants - unlike most other ants - face severe time constraints when generating new foraging routes each day. While ants living in a permanent nest site may thoroughly explore their environment [20, 21] or clear trails of vegetation [22, 23, 24, 25] in order to create relatively straight and efficient foraging paths, army ant trails often weave tortuously through the complex tropical forest floor [26, 19].

In order to improve the efficiency of their trails, army ants are capable of linking their own bodies together to dynamically create physical structures along the foraging path [18, 19, 27, 28, 16]. These structures may be used to widen paths to increase the flux of ants, or to form bridges over gaps in the terrain (reaching spans of over 12 cm, or approximately 12 ant body lengths) to decrease the tortuosity of their trails [29, 16]. Ants modulate their bridge-building behavior in response to local information, allowing these bridges to adapt to current traffic conditions, recover from damage, and dissemble when underused [28], so that they exist as needed at particular points along the trail.

While the living bridges can increase the flow of ants and resources along trails, they also impose a cost on the colony. The ants forming the bridges are locked into the structure, sometimes for several minutes at a time [28], preventing them from participating in other important foraging activities such as capturing and killing prey or transporting food items along the trail. Understanding the overall effect of these living structures on the colony's foraging rate requires a quantification of both the benefit (here, shortening the travel distance) and cost (here, removing ants from the foraging pool) of each structure, and converting these to the common currency of overall foraging rate.

In a recent study, Reid et al. [16] experimentally manipulated living bridges built by colonies of *E. hamatum* and measured these benefits and costs. The researchers inserted deviations into an existing foraging trail (Figure 1) and recorded the formation of bridges on the experimental apparatus. They showed that bridges typically initiated at the bend of the deviation but over time grew and moved away from the initiation point to increasingly shortcut the deviation. However, the final, steady state, position of the bridges tended to not fully minimize the trail length. Instead, the distance that the bridge traveled depended on the angle of the apparatus deviation [16].

**Figure 1:**
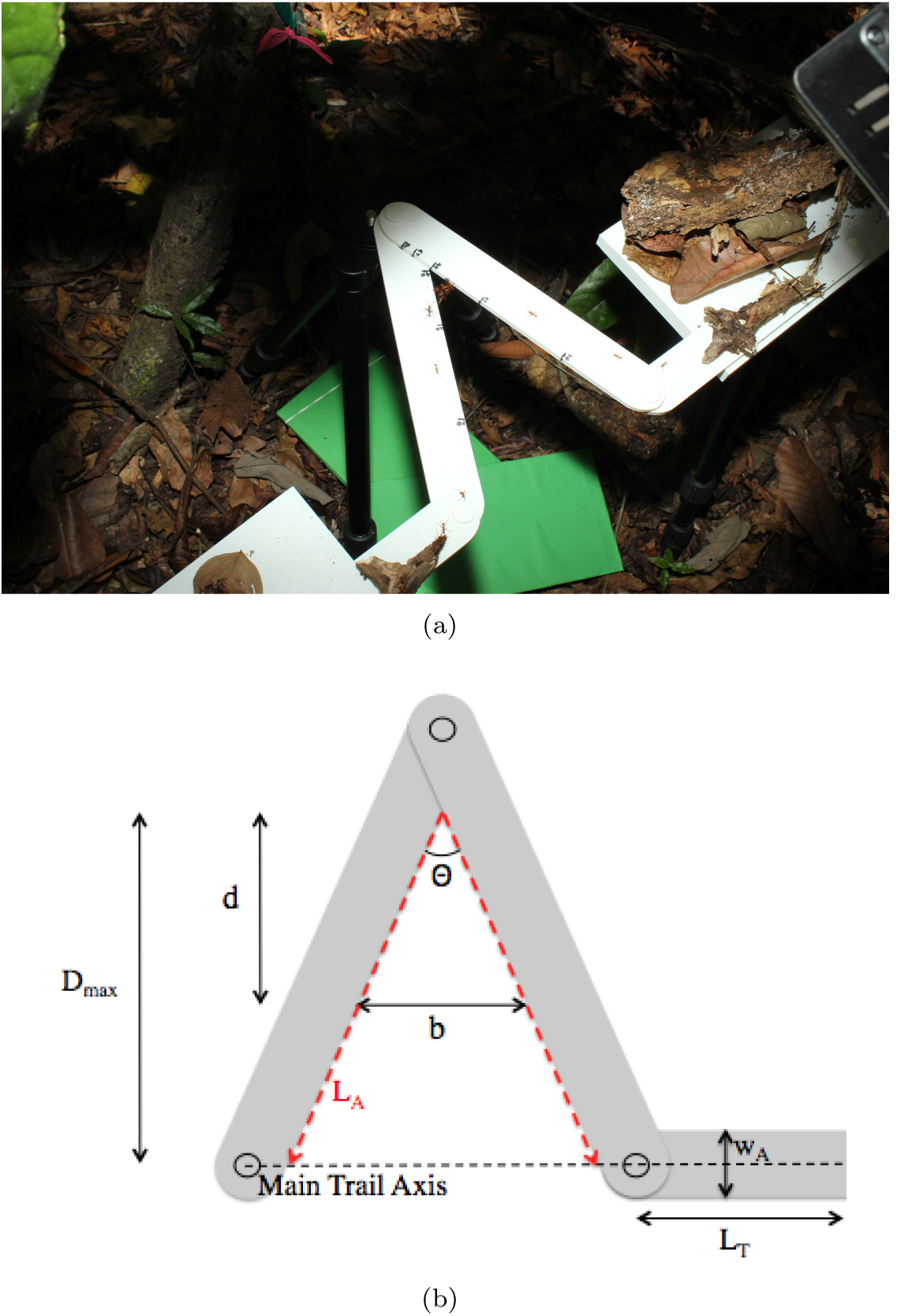
1(a) Field apparatus used in Reid et al. [16] to experimentally manipulate living bridges built by colonies of army ant *Eciton hamatum.* 1(b) Schematic representation of the experimental apparatus introduced into a foraging trail of army ants in Reid et al. [16]. The introduction of such an apparatus has the effect of adding an additional length *L_A_* to the distance (*L_T_*) foraging ants must travel. In order to short-cut this additional distance, army ants construct a living bridge that initially forms at an apex of angle *θ*, and moves down toward the main trail axis until reaching some optimal position. Here *w_A_* is the width of an apparatus arm.

Reid et al. [16] also counted the number of ants required to build a bridge of a certain length. Crucially, the bridges were observed to widen as they lengthened, so that while the travel distance saved increased linearly with bridge length, the number of ants diverted from the foraging pool increased quadratically with bridge length. Changes in travel distance and the number of available foraging ants can be converted directly to changes in the density of foraging ants on the trail, which the researchers used as a proxy for foraging rate. Maximizing the foraging rate as a function of bridge position led to a unique, non-trivial, optimal position, which the researchers showed could be matched closely to the empirically observed bridge positions.

Here, we propose to extend the mathematical model introduced by Reid et al. [16] into a much broader theoretical framework. This framework can be used to derive optimization models for bridge formation patterns that occur in response to a much wider range of geometrical scenarios along a trail. We demonstrate this framework for two new classes of scenarios. First, we explore asymmetric deviations along the foraging trail, for which the optimal bridges, unlike those observed in Reid et al. [16], are predicted to be generally not parallel to the main foraging trail. For this scenario, we also demonstrate the optimal formation process, which optimizes foraging rate throughout the duration of bridge construction. Second, we study scenarios with multiple consecutive deviations, which we predict will lead to multiple bridges. In such scenarios, the position of one bridge constrains the possible positions of the other bridges, so that the geometry forces more complex coordination in optimal bridge positioning. Our results produce quantitative predictions about the number of bridges, their position relative to the apparatus (or natural obstacle) and to other bridges, and their angle relative to the main foraging trail, which can be directly tested in field experiments. In addition to allowing us to derive some specific predictions, consideration of these two classes of scenarios simultaneously shows how to adapt the modeling paradigm used here for additional scenarios not considered in this work.

## 2. Methods

### 2.1. General model framework

As in Reid et al. 16, we consider a foraging trail of total length *L_T_* and N army ants. If an experimental apparatus (or natural obstacle) is introduced, then this adds an additional length of *L_A_* to the distance over which the ants must travel (Figure 1). Thus, in the absence of any living bridges the overall foraging density of ants is

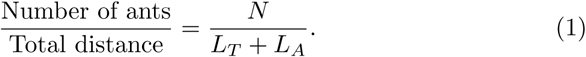

The additional length *L_A_* added to the travel path by the placement of a particular apparatus within the trail depends on the geometric configuration of the apparatus, due to the physical implementation of the experimental set-up, as well as our assumption that ant trails tend to follow the edge of the apparatus in order to minimize travel distance (see Reid et al. [16]). We estimate the number of ants N by multiplying the average empirically measured density of foraging ants by *L_T_* + *L_A_*, as in Reid et al. [16].

The presence of one (or more) bridges will modify both the number of ants moving on the trail (since the bridges are themselves comprised of ants) and the distance of travel, both of which will modulate the traffic density on the trail. The number of available foraging ants becomes N minus the number of ants sequestered in the bridge structures, *i.e.*

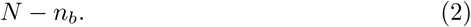

However, there are substantial functional differences between a typical bridge-building and non-bridge-building (i.e., foraging) ant. Ants that take up positions in bridges tend to be smaller and less effective at capturing and carrying prey items [30, 31]. Therefore, if our unit of ants is assumed to be foraging ants, then the cost of including a (smaller) ant into the bridge structure will actually be less than one foraging ant. In order to account for these size and functional differences between bridge-building and foraging ants, we introduce a free parameter *α* in order to make direct comparisons between model results and experimental data. Thus we modify equation (2) to become

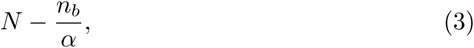

where we set *α* = 17.02 as in Reid et al. [16], but which may need to be refit when testing new ecological conditions (such as nighttime colony migrations, where the functional differences between ants of different sizes may not be the same as when foraging).

In the presence of bridges, the distance of travel, *f*, becomes

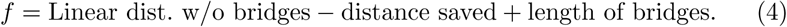

Thus, in the presence of bridges, the density function to be optimized is

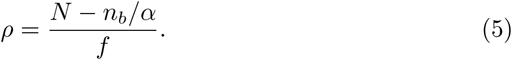

In order to maximize the density of foraging ants for a particular geometric configuration of apparatuses (or obstacles in nature), all that is left is to describe how the number of ants in the bridge *n_b_* and trail length *f* vary with bridge position.

### 2.2 Asymmetric scenario

#### 2.2.1. Optimal bridge position

While the experimental apparatus of Reid et al. [16] was symmetric relative to the main trail axis (Figure 1), in nature obstacles will generally be asymmetric. This may yield optimal bridges that are at a nonzero angle with the main trail axis, unlike the parallel bridges observed in Reid et al. [16]. In our first extended model, we allow for a difference in the angle that the left (*θ*) and right (*ϕ*) arms of the apparatus respectively make with the line perpendicular to the main trail axis (Figure 2). Furthermore, we allow for a difference in the lengths of each arm of the apparatus.

**Figure 2:**
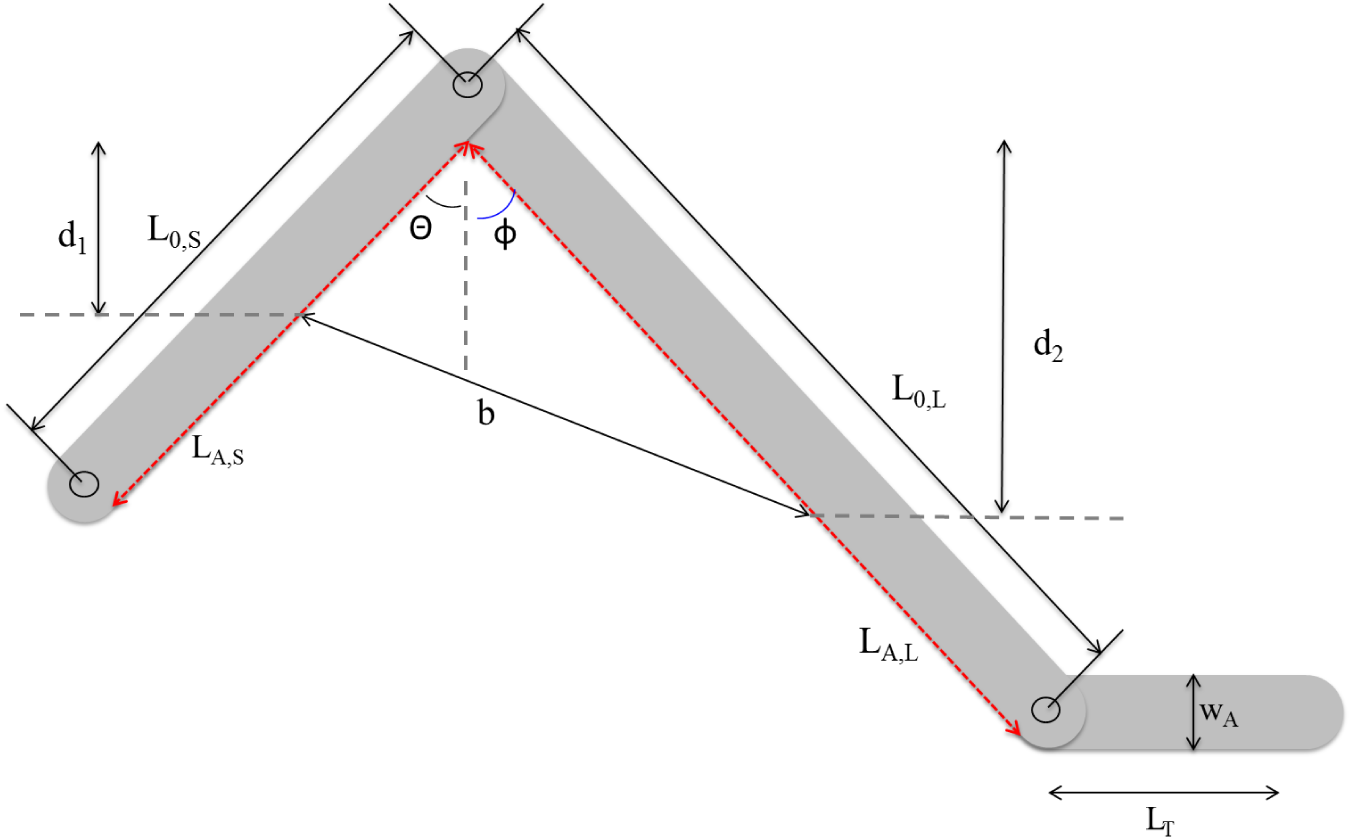
Schematic of a theoretical apparatus that is predicted to result in the construction of army ant living bridges that are not necessarily parallel to the main trail axis. The additional length of such an apparatus added to the path of travel is *L_A,S_* + *L_A,L_*. Here *w_A_* is the width of an apparatus arm. All other variables and parameters are described in tables 1 and 2.

Let *L*_0_,*s* and *L*_0_,_*L*_ be the hinge-to-hinge distance along the left and right arms, respectively, of the apparatus. Then the lengths along each arm from the apex of the apparatus to the unconnected end of each arm is

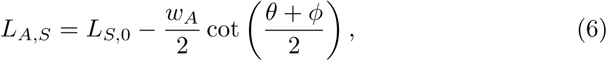

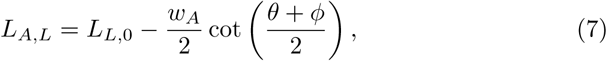

where *w_A_* is the width of the apparatus arm. Thus, *L_A_=L_A,S_+L_A,L_,* and the maximum possible vertical distances that a living bridge could travel relative to the left and right arms respectively is

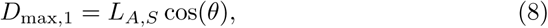

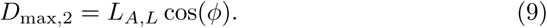

In this case, the linear length of a living bridge formed to cross from the left arm to the right, according to the law of cosines, will be

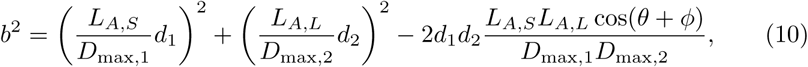

where each of the parameters and variables is listed and described in Tables 1 and 2. Furthermore, the linear distance of travel for the ants is now

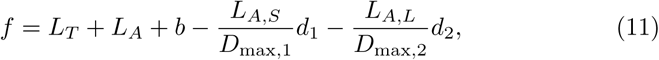

and we obtain our density function

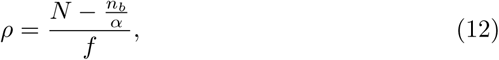

where the number of ants sequestered for the formation of a bridge is

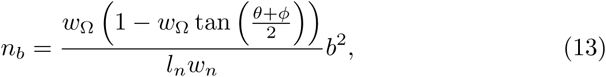

again with all relevant parameters listed and described in Tables 1 and 2. This equation for the number of sequestered ants takes into account the length (*l_n_*) and width (*w_n_*) of a typical bridge ant, as well as the scaling *w*_Ω_ between the width and length of a living bridge (see Reid et al. [16]).

**Table 1:** Fixed parameters used in all models.

| Notation | Description | Value | Units |
| --- | --- | --- | --- |
| $L_T$ | trail length without bridges | 100 | cm |
| $l_n$ | length of an average ant when occupying a position within the bridge structure | 0.691 | cm |
| $w_n$ | width of an average ant when occupying a position within the bridge structure | 0.107 | cm |
| $w_A$ | width of apparatus arm | 3.3 | cm |
| $\alpha$ | free parameter to adjust the space occupied by an ant on the trail | 17.02 | |

**Table 2:** Geometric parameters and variables corresponding to asymmetric model representing an apparatus such as shown in Figure 2.

| Notation | Description | Value | Units |
| --- | --- | --- | --- |
| $L_{S,0}$ | Hinge-to-hinge length of left apparatus arm | 22 | cm |
| $L_{L,0}$ | Hinge-to-hinge length of right apparatus arm | 44 | cm |
| $\theta, \phi$ | Angle of left and right arm respectively from the vertical | 0 to 45 | degrees |
| $w_\Omega$ | Ratio between width and length of a bridge, value from Reid et al. [16] | $4.799(\theta + \phi)^{-0.5014}$ | N/A |
| $L_{A,S}$ | Travel length along left arm from apex to opposite hinge | $L_{S,0} - \frac{w_A}{2} \cot\left(\frac{\theta+\phi}{2}\right)$ | cm |
| $L_{A,L}$ | Travel length along right arm from apex to opposite hinge | $L_{L,0} - \frac{w_A}{2} \cot\left(\frac{\theta+\phi}{2}\right)$ | cm |
| $L_A$ | Sum of travel lengths along each arm | $L_{A,S} + L_{A,L}$ | cm |
| $D_{\max,1}$ | Maximum vertical distance from apex to bottom of left arm | $L_{A,S} \cos(\theta)$ | cm |
| $D_{\max,2}$ | Maximum vertical distance from apex to bottom of right arm | $L_{A,L} \cos(\phi)$ | cm |
| $d_1$ | Vertical distance of bridge from apex to position on left arm | 0 to $D_{\max,1}$ | cm |
| $d_2$ | Vertical distance of bridge from apex to position on right arm | 0 to $D_{\max,2}$ | cm |

In order to determine the optimal bridge position, we optimize the density as a function of *d*_1_ and *d*_2_ for a range of fixed values of *θ* and *ϕ*. Since it is difficult to reliably solve for the maxima of (12) analytically, the optimization is carried out via numerical routines using the R package DEoptim [32, 33, 34, 35, 36]. We note that the density function described by equation (12) is continuous and is to be maximized over the rectangle [0, *D*_max,1_] × [0, *D*_max,2_]. This guarantees the existence of a maximum value. The results are presented in the next section. As will be seen, an interesting consequence of asymmetry is that there may be transitions in the orientation of bridges that are formed at an angle to the main trail axis.

#### 2.2.2. Optimal bridge formation process

We also derive the optimal bridge formation process by asking, for a fixed bridge length, what the position is that maximizes foraging rate. We then vary the bridge length from 0 to the final optimal length and calculate the optimal bridge position for each length, thereby constructing the optimal bridge-building trajectory that maximizes foraging rate for the entire duration of the bridge formation process.

Let *b_f_* represent the bridge length parameter. When *b_f_* is fixed, through equations (10)-(12) we arrive at a constrained optimization problem. That is, we seek to optimize the density

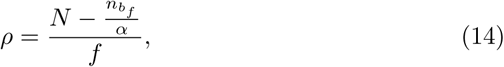

subject to the constraint

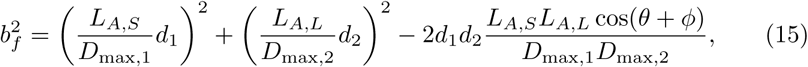

where

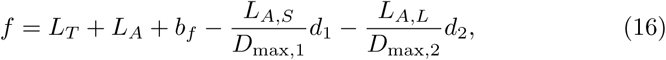

and

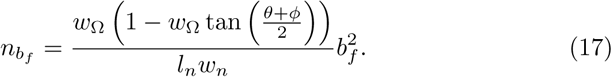

We note that *n_bf_* and hence 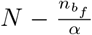 are now held constant since *b_f_* is fixed.

The values for *L_A_*_,*S*_, L_*A,L*_, *D*_max,1_ and *D*_max, 2_ are obtained just as before.

In the appendix, we derive the values of *d*_1_ and *d*_2_ in the intervals [0, *D*_max,1_] and [0, *D*_max, 2_] respectively that maximize equation (14) subject to the constraint given by equation (15).

### 2.3. Multiple obstacles scenario

In addition to asymmetric obstacles, in nature there may frequently be multiple consecutive obstacles, leading to multiple deviations from a straight trail (e.g., Figures 3 and 4). In these scenarios, more than one bridge may form in sequence, and the position of one bridge may constrain the possible positions of downstream bridges. Thus, this introduces a coupling between bridges that must be accounted for in the optimization process. In our second model, we consider multiple, but symmetric, deviations from the trail, which we predict to produce more than one bridge that coordinate in a striking fashion. Figures 3 and 4 illustrate apparatuses that could be expected to lead to the formation of two and three bridges, respectively.

**Figure 3:**
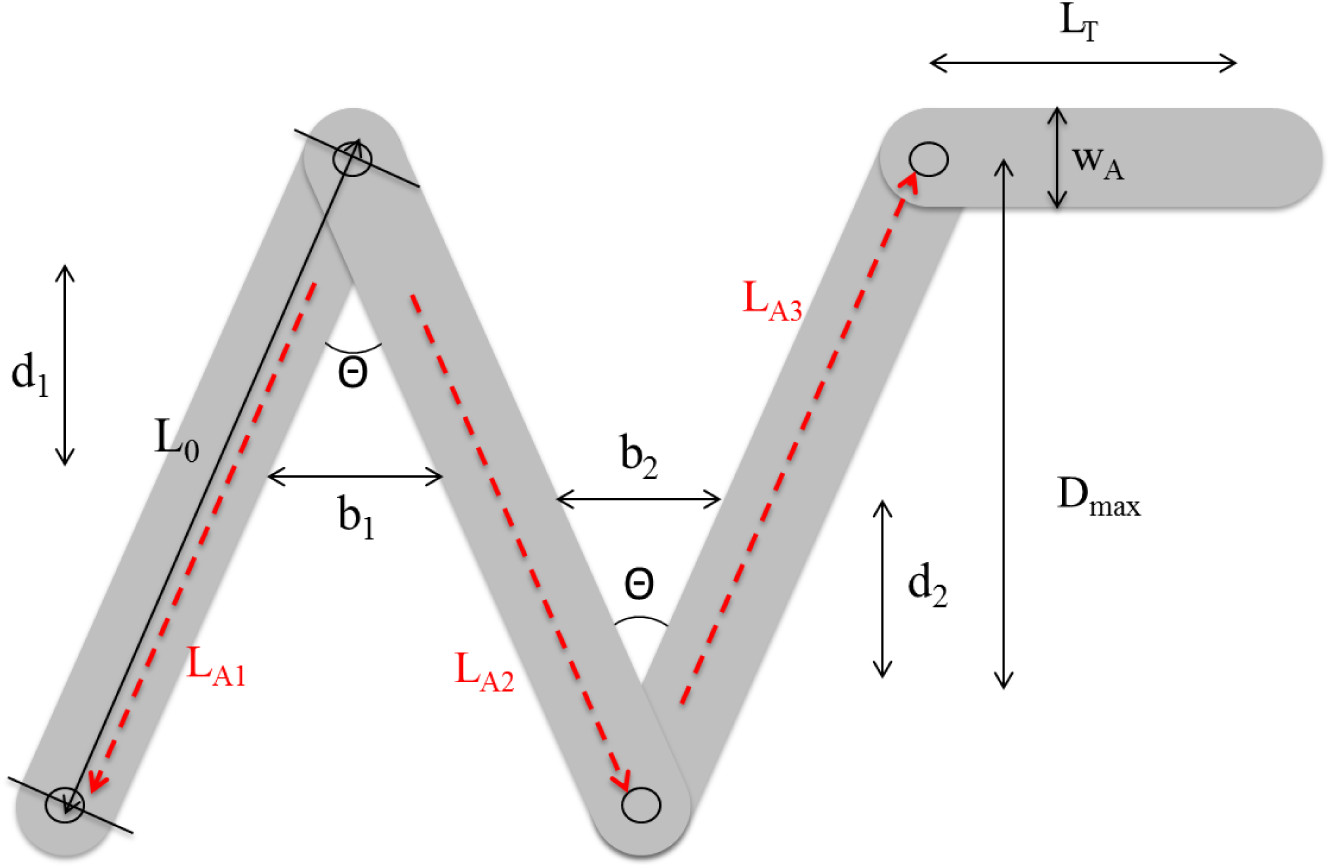
Schematic of a theoretical apparatus that is predicted to result in the construction of two distinct army ant living bridges. The additional linear length of such an apparatus added to the path of travel is well-approximated by *L_A_* = *L*_*A*_1__ + *L*_*A*_1__ + *L*_*A*_1__. Note that, due to symmetry *L*_*A*_1__ = *L*_*A*_2__ = *L*_*A*_3__. Here *w_A_* is the width of an apparatus arm. All other variables and parameters are described in tables 1 and 3.

**Figure 4:**
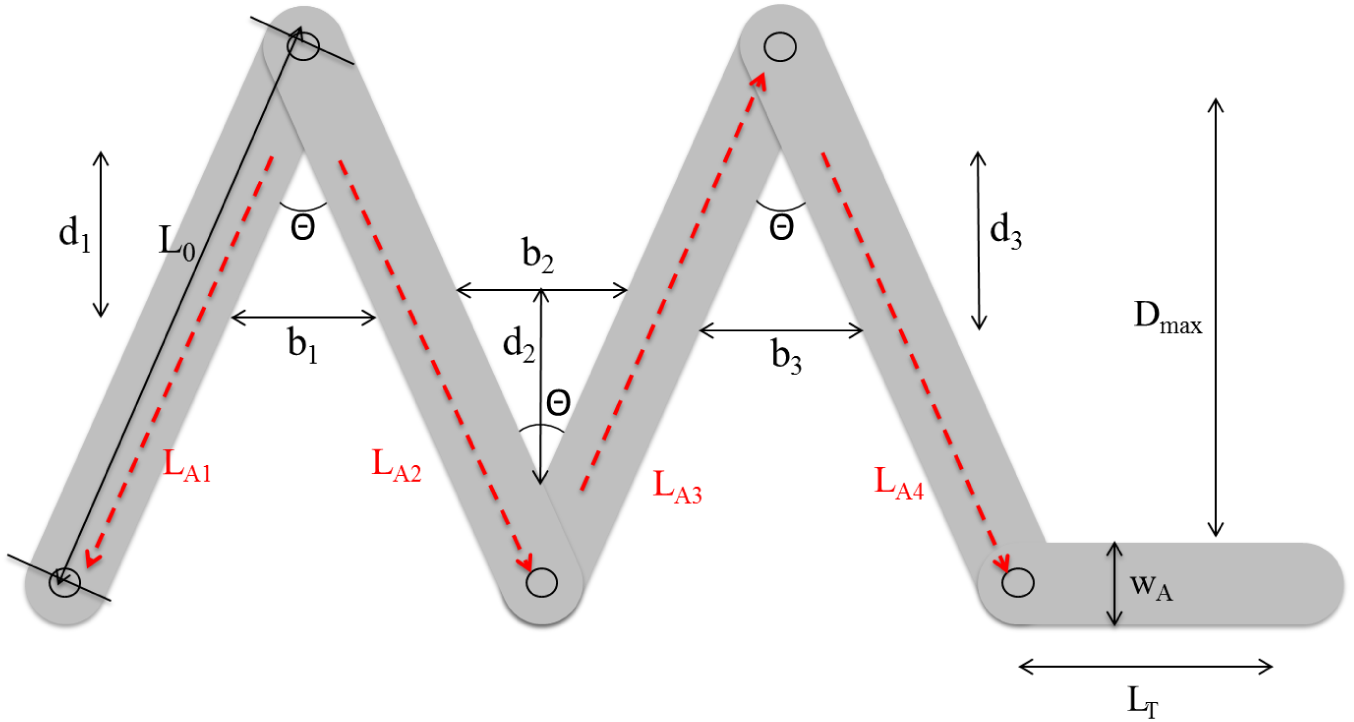
Schematic of a theoretical apparatus that is predicted to result in the construction of three distinct army ant living bridges. The additional linear length of such an apparatus added to the path of travel is well-approximated by *L_A_* = *L*_*A*_1__ + *L*_*A*_2__ + *L*_*A*_3__ + *L*_*A*_4__. Note that, due to symmetry, *L*_*A*_1__ = *L*_*A*_2__ = *L*_*A*_3__ = *L*_*A*_4__. Here *w_A_* is the width of an apparatus arm. All other variables and parameters are described in tables 1 and 4.

For the two bridge case illustrated in Figure 3, let *L*_0_ be the hinge-to-hinge length along each of the three segments of the apparatus. Then we have that the distance from the apex along each arm is

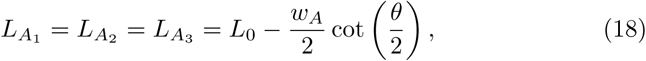

so *L_A_* = *L*_*A*_1__ + *L*_*A*_2__ + *L*_*A*_3__ = 3*L*_*A*_1__, and the maximum vertical distance each bridge can travel is

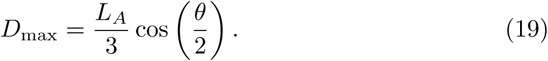

Then, our density function will take the form

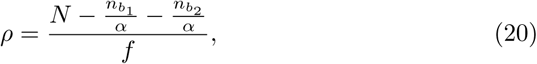

where the number of ants sequestered in each bridge is

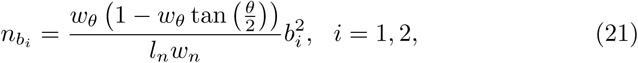

and the length of each bridge is

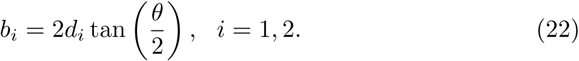

Then, the linear distance of a trail with two bridges will be

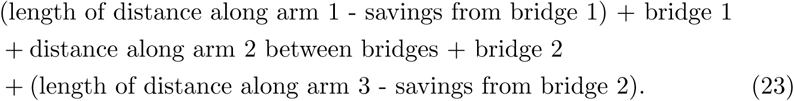

Thus, we have

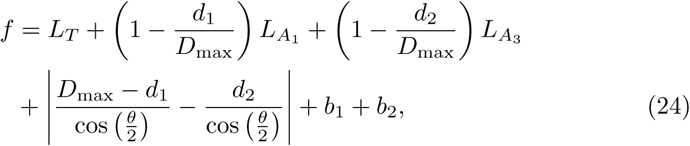

where all of the parameters are listed and described in Tables 1 and 3.

**Table 3:** Geometric parameters and variables corresponding to two-apex apparatus such as shown in Figure 3.

| Notation | Description | Value | Units |
| --- | --- | --- | --- |
| $L_0$ | Hinge-to-hinge length of each arm | 22 | cm |
| $\theta$ | Angle of each apex | 0 to 60 | degrees |
| $w_\theta$ | Ratio between width and length of a bridge, value from Reid et al. [16] | $4.799\theta^{-0.5014}$ | N/A |
| $L_{A_1}, L_{A_2}, L_{A_3}$ | Travel length along each arm of apparatus from apex to opposite hinge | $L_0 - \frac{w_A}{2} \cot\left(\frac{\theta}{2}\right)$ | cm |
| $L_A$ | Sum of travel lengths along each arm | $L_{A_1} + L_{A_2} + L_{A_3}$ | cm |
| $D_{\max}$ | Maximum vertical distance from apex to end of each arm | $\frac{L_A}{3} \cos\left(\frac{\theta}{2}\right)$ | cm |
| $d_1$ | Vertical distance of bridge from apex to position on arms forming first apex | 0 to $D_{\max}$ | cm |
| $d_2$ | Vertical distance of bridge from apex to position on arms forming second apex | 0 to $D_{\max}$ | cm |

Putting the last four equations together gives a complete expression for the density as a function of the variables *d*_1_ and *d*_2_, which when optimized provides the best positioning of the two bridges to maximize the density of foraging ants on the trail. Again, the optimization is carried out via the DEoptim package in R, and the results are presented in the next section.

A straightforward extension of the two bridge case leads to the following model for a situation such as illustrated in Figure 4, that may lead to the formation of three bridges. In this case we obtain

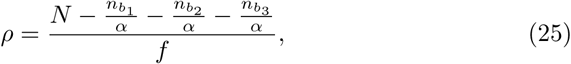

with

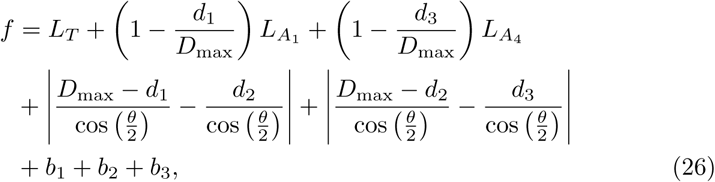

and with the bridge lengths and the number of ants sequestered for bridge formation given by the same expressions as in the two bridge case. This model can easily be extended to arbitrary numbers of obstacles, and can be generalized to multiple asymmetric obstacles. A term of the form

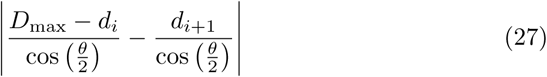

such as appears in equations (24) and (26) is called the **mid-distance** between two consecutive bridges.

We note that each of the density functions in equations (12), (20) and (25) is guaranteed to have a biologically relevant maximum value since it is a continuous function maximized over a closed and bounded set of possible distance values. Also, in each of the equations (12), (20) and (25), the number of ants sequestered for bridge formation depends quadratically on the distance(s), while the linear distance of travel *f* depends on the distance(s) in such a way as to be of order to the first power. The consequence of this observation for the cost-benefit trade-off of living bridge formation is already pointed out in Reid et al. [16]. However, unlike in Reid et al. [16], the presence of square roots, *e.g.* in (11), and absolute values, *e.g.* in (26), make the equations much less tractable to solving analytically for exact expression for optimal bridge-positioning. Additional discussion of this and some related points are provided in the appendix.

### 2.4. Model parameters

In order to implement the models derived from the theory in the previous subsections, we choose parameter values that are either taken directly from Reid et al. [16], or chosen to be consistent with values from Reid et al. [16]. Specifically, for the lengths of the sides of a proposed or theoretical apparatus, we choose parameter values that are on the order of the lengths of the experimental apparatus from Reid et al. [16]. The values for all parameters used to obtain the following results are listed in Tables 1-4. Note, however, that the theory does not depend on the explicit values of the geometric parameters. Therefore, our approach can be adapted to the specifics of an experimental apparatus or a naturally occurring obstacle, provided that it can be described with a similar set of geometric parameters that completely specifies its configuration.

**Table 4:** Geometric parameters and variables corresponding to three-ape apparatus such as shown in Figure 4.

| Notation | Description | Value | Units |
| --- | --- | --- | --- |
| $L_0$ | Hinge-to-hinge length of each arm | 22 | cm |
| $\theta$ | Angle of each apex | 0 to 60 | degrees |
| $w_\theta$ | Ratio between width and length of a bridge, value from Reid et al. [16] | $4.799\theta^{-0.5014}$ | N/A |
| $L_{A_1}, L_{A_2}, L_{A_3}, L_{A_4}$ | Travel length along each arm of apparatus from apex to opposite hinge | $L_0 - \frac{w_A}{2} \cot\left(\frac{\theta}{2}\right)$ | cm |
| $L_A$ | Sum of travel lengths along each arm | $L_{A_1} + L_{A_2} + L_{A_3} + L_{A_4}$ | cm |
| $D_{\max}$ | Maximum vertical distance from apex to end of each arm | $\frac{L_A}{4} \cos\left(\frac{\theta}{2}\right)$ | cm |
| $d_1$ | Vertical distance of bridge from apex to position on arms forming first apex | 0 to $D_{\max}$ | cm |
| $d_2$ | Vertical distance of bridge from apex to position on arms forming second apex | 0 to $D_{\max}$ | cm |
| $d_3$ | Vertical distance of bridge from apex to position on arms forming third apex | 0 to $D_{\max}$ | cm |

### 2.5. Model extensions

As part of supplementary material, we have developed freely available code using the R programming language, R Core Team [37], that can be used to implement the models from this paper or any similar models that one may derive. This package can be found at https://goo.gl/zam27s.

The derivations for the density functions given by equations (12), (20) and (25) illustrate how an appropriate density function may be derived for any number of different configurations, and for an apparatus or obstacle that an army ant foraging trail may encounter that could result in the formation of one or more living bridges. This work serves to show that we have a general theory that can be used to predict army ant living bridge positioning for a variety of actual and conceivable scenarios. Further, the theory may also be coupled with agent-based simulations or other dynamic modeling approaches to provide even more detailed computational studies of living bridge formation, such as the individual behavioral rules that may lead to the dynamic formation of such optimal structures. In the next section, we present the quantitative and qualitative results predicted by our theory for the three specific configurations captured by equations (12), (20) and (25).

## 3. Results and Discussion

### 3.1. Asymmetric scenario

In general, optimal bridges in asymmetric apparatuses are not parallel to the main trail axis. Examples of optimal bridge positions are illustrated in Figure 5(a)-5(b). To obtain these results, we fixed one of the arm angles, *θ,* to be either 20° or 10°, and varied the other angle, *ϕ*, from 5° to 30°. We then calculated the optimal bridge position for each combination of angles as the maximizing distances^2^, *d*_1_ and *d*_2_, that the two ends of the optimal bridge travel down each arm of the apparatus from the apex (Figure 2), and plotted the the difference between these two lengths. Here, a negative difference indicates that the optimal bridge travels further down the arm associated with the angle *ϕ*, whereas a positive difference indicates that the bridge travels further down the arm associated with the angle *θ*.

**Figure 5:**
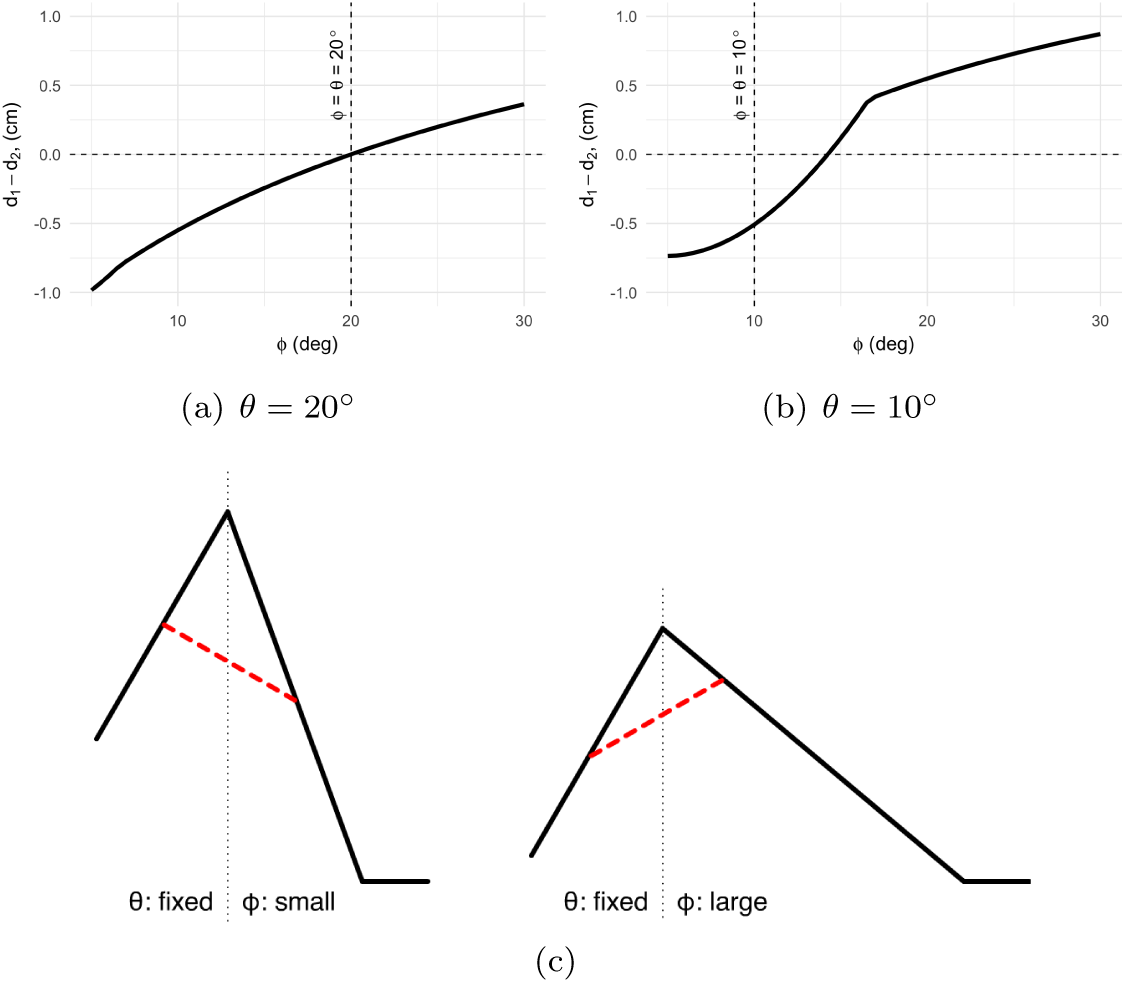
Quantitative (5(a) - 5(b)) and qualitative (5(c)) results described in section 3.1. These results show the predicted arrangement of optimal living bridge configurations for an asymmetric apparatus as a function of the angle *ϕ* with the angle *θ* fixed at 20° (5(a)) and 10° (5(b)) respectively. Equation (60) implies that a living bridge forms at the apex of the apparatus and quickly establishes an angle with respect to the main trail axis that is completely determined by the ratio 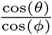 This angle with respect to the main trail remains constant as the living bridge moves down the apparatus to its equilibrium position, at least until the bridge “runs out of road” along one arm or the other. That is, for fixed *θ* and *ϕ*, the orientation of the living bridge only changes as the bridge moves down the apparatus if it reaches the bottom of the shorter side before establishing its equilibrium position. Computed for a foraging density of approximately 2.2.

Typically, the optimal bridge travels further down the apparatus arm with a smaller angle (Figure 5(a)). This is shown by the difference in bridge end positions being negative when *ϕ* < *θ* and positive when *ϕ* > *θ*. Specifically, we find as derived in the appendix that

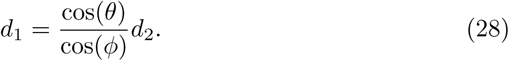

This relationship implies that when the two angles *ϕ* and *θ* are equal, the optimal bridge tends to be parallel to the main trail axis. Therefore, the symmetric experimental apparatus studied in Reid et al. [16] is shown to be a special case of the more general asymmetric apparatus.

However, when the angle *θ* is small, we observe deviations from the above behavior. In this case, bridges parallel to the main trail axis can form even when the two angles are not equal (Figure 5(b)). When the two angles are equal, the bridge is skewed such that it is further down the arm associated with the angle *ϕ*. This is due to the unequal lengths of the two arms of the apparatus (Figure 2). The arm associated with the angle θ is shorter than that associated with angle *ϕ*, and the arm lengths define the maximum distance that the bridge can travel. In this regime, one end of the bridge meets the maximum distance of the shorter arm, but the other end continues to travel further down the longer arm. This phenomenon also explains the presence of a 'kink' in the curve in Figure 5(b).

Nonetheless, the trend that the difference in distance increases as the angle *ϕ* increases is still observed, such that a similar transition from a negative difference to a positive difference occurs, albeit when *ϕ* > *θ*. Therefore, in general we predict bridges to travel further down the arm associated with angle *ϕ* when *ϕ* is small, and travel further down the arm associated with angle *θ* when *ϕ* is large. This prediction holds as parameters are varied and for a wide range of angles.

We can predict not only the optimal final position of a living bridge, but also the optimal bridge formation process, by constraining the bridge length to certain values shorter than the final length and solving for the optimal bridge at each length (the appendix contains the details of solving this constrained optimization problem). Because equation (28) is true for any bridge length, it predicts that a living bridge forms at the apex of the apparatus and quickly establishes an angle with respect to the main trail axis that is completely determined by the ratio 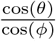. This angle with respect to the main trail remains constant as the living bridge moves down the apparatus to its equilibrium position, at least until the bridge “runs out of road” along one arm or the other. That is, for fixed *θ* and *ϕ*, the orientation of the living bridge only changes as the bridge moves down the apparatus if it reaches the bottom of the shorter side before establishing its equilibrium position.

### 3.2. Multiple obstacles scenario

When there are multiple obstacles, we find that multiple bridges form, although, as with asymmetric obstacles, the position of the bridges depends on the angle of the apparatus (Figures 6 and 7. For the case of two obstacles (6), small and moderate angles result in optimal bridges which are situated halfway between the two apexes, such that the two bridges form a straight path. In this regime, as the angle of the apparatus increases, the maximal possible distance that a bridge can move from the apex, *D*_max_, increases, and both bridges have distance 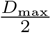 from each apex (note that *D*_max_ increases, rather than decreases, with the apparatus angle because of the nonzero width of the apparatus arm, see Methods and [16]). Furthermore, the distance between the two bridges remains 0. At large angles, however, the bridges are predicted to separate and move closer to their respective apex as the angle increases further. This is illustrated by the distance of the two bridges from the apex decreasing and the distance between the two bridges increasing.

**Figure 6:**
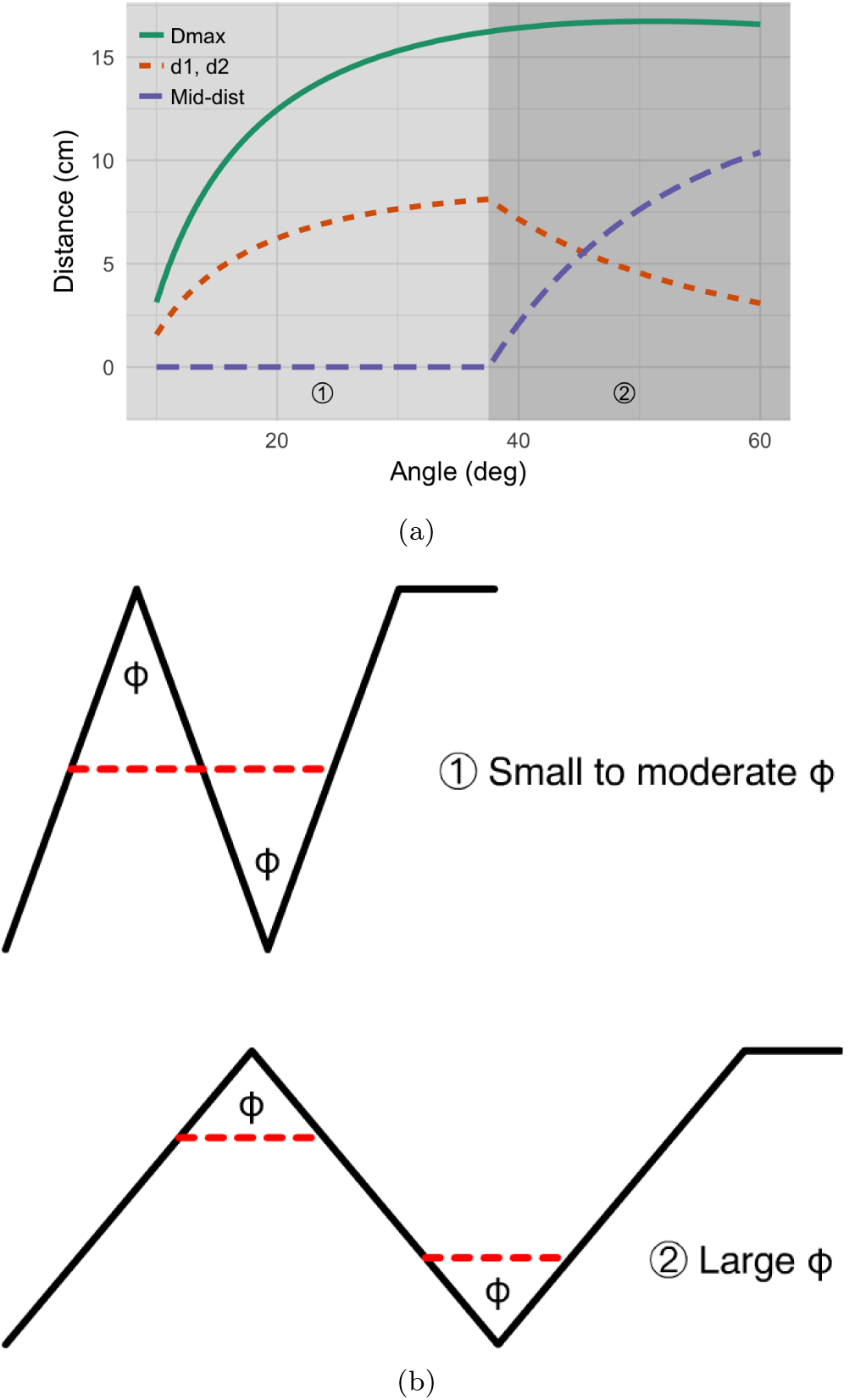
Figure 6(a) shows, as a function of angle *θ*, the greatest possible vertical distance a living bridge can travel along an apparatus, the distance values *d*_1_ and *d*_2_ that optimize the density function (20) and determine the optimal positioning of a living bridge, and the linear distance between two bridges at their optimal positioning. Thus, the theory predicts that optimal bridge formation for an apparatus such as Figure 3 is such that the linear distance between two bridges along a common arm is minimized as much as the availability of bridgebuilding ants allows for under the given geometric constraints imposed by apex angle. For the quantitative results presented here, a foraging density value of approximately 2.2 is used. Qualitatively similar results are obtained for a variety of different parameter values.

**Figure 7:**
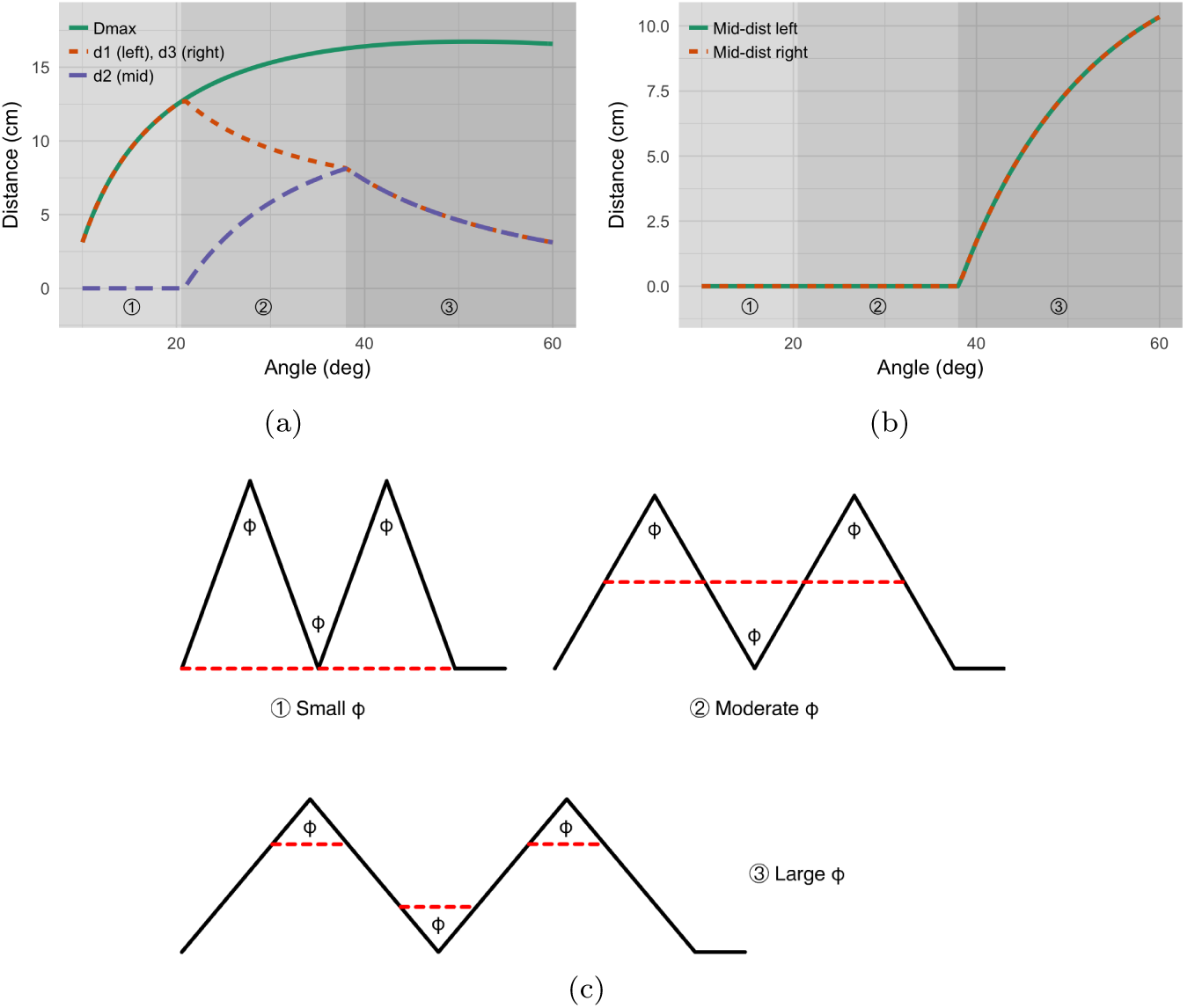
Figure 7(a) shows, as a function of angle θ, the greatest possible vertical distance a living bridge can travel along an apparatus and the distance values *d*_1_, *d*_2_, and *d*_3_ that optimize the density function (25) and determine the optimal positioning of a living bridge. Figure 7(b) shows the linear distance between two consecutive bridges at their optimal positioning. Thus, the theory predicts that optimal bridge formation for an apparatus such as Figure 4 is such that the linear distance between all three bridges along common arms is minimized as much as the availability of bridge-building ants allows for under the given geometric constraints imposed by apex angle. For the quantitative results presented here, a foraging density value of approximately 2.2 is used. Qualitatively similar results are obtained for a variety of different parameter values.

For the case of three obstacles, there are three, rather than two regimes (Figure 7). For small apparatus angles, only two bridges form, which together extend the main trail axis in a straight line. Here, as the angle increases, the maximum possible distance from the apexes (*D*_max_) increases, and the outer bridges remain at the maximum distance *d*_1_ = *d*_2_ = *D*_max_, while the inner bridge has zero length. For moderate angles, there are three bridges, which together form a straight line. As the angle increases in this regime, the bridges move increasingly toward the middle of the apparatus. At large angles, the three bridges separate, as in the two-obstacle case, and all three bridges have equal length. As the apparatus angle increases further, each bridge moves towards its respective apex.

### 3.3 The role of ant density

In nature, the density of ants on a trail can vary dramatically, depending on the time of day, the size of the colony, and the amount of available food in the trail’s vicinity. We investigated how ant density affects our previous results. We show results for the asymmetric apparatus, although we find qualitatively similar results for the two-obstacle and three-obstacle scenarios.

In general, the effects of changing the angle of the apparatus become larger as ant density increases (Figures 8(a) and 8(b)). However, the qualitative features of our results remain the same across densities. Thus, performing experiments on trails with higher densities of ants will improve the ability to detect the patterns of bridge formation that we predict, if ants build bridges in order to maximize foraging rate as hypothesized in Reid et al. [16].

**Figure 8:**
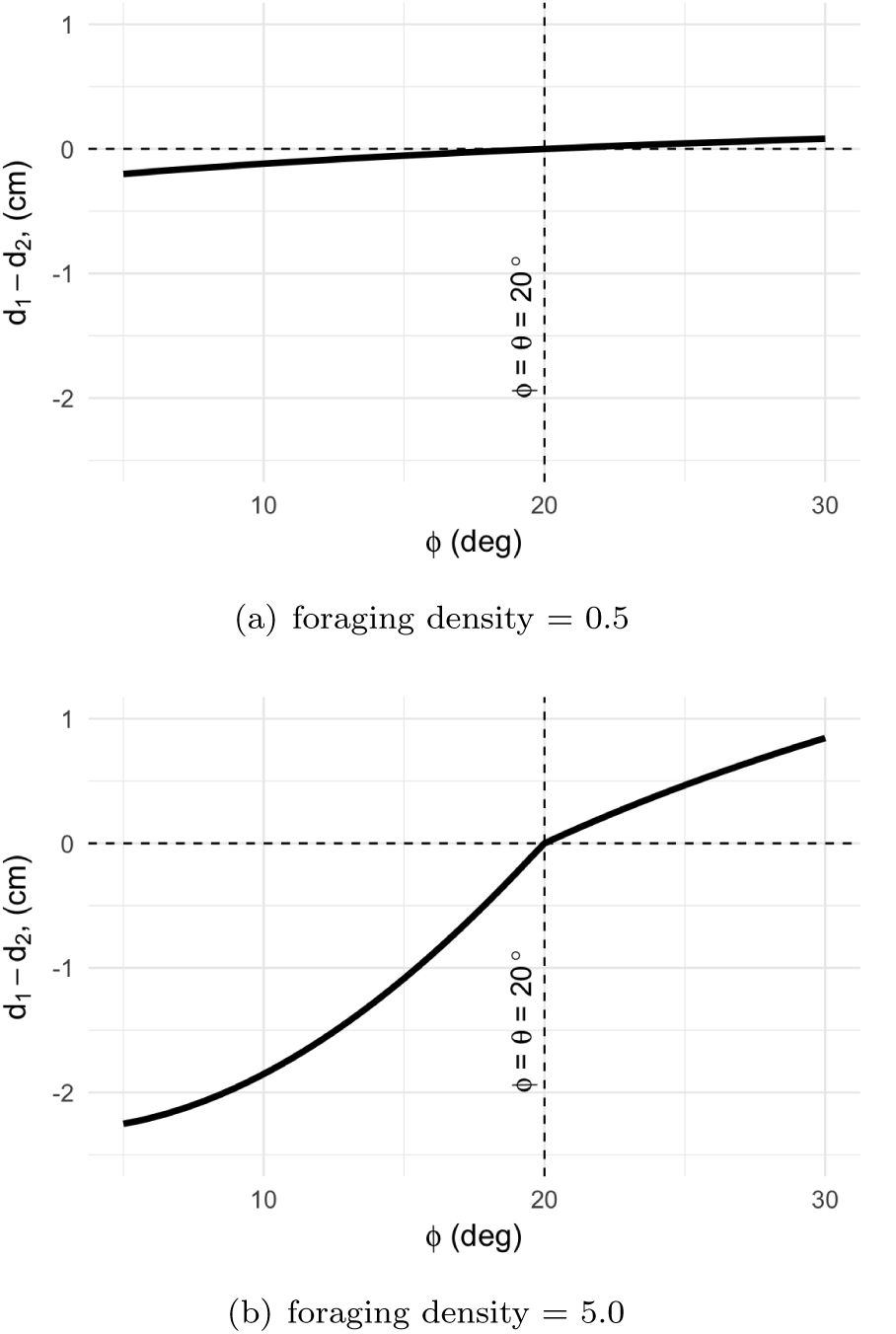
In nature, the density of ants on a trail can vary dramatically. Motivated by this, we investigated how ant density affects our previous results. For example, we compare the quantitative results as shown in figure 5(a) with those obtained by decreasing (8(a)) and increasing (8(b)) the density. In general, the effects of changing the angle of the apparatus become more larger as ant density increases. However, the qualitative features of our results remain the same across densities. The situation is highly similar for the other apparatus configurations considered in this work. Thus, the predictions made by the theory are robust with respective to the qualitative behavior predicted.

## 4. Conclusion

Detemining the details of the construction of army ant living bridges is important to understanding the collective behavior of army ants. We extended a mathematical model for a specific case living bridge construction into a broad theoretical framework that may be applied to a variety of increasingly complex natural and experimental obstacles, which are predicted to result in the formation of a living bridge by foraging army ants. Using this framework, we made explicit predictions that can be experimentally tested. In particular, for each scenario, we identified qualitatively different bridge-building regimes, depending on the configuration of the experimental apparatus, which will be more amenable to testing in the field. If the living bridges that army ants construct function mainly to maximize foraging rate, then these different regimes will be observed in nature.

## Appendix

In this appendix, we expound additional properties of the theory presented in the main body of this work by carrying out a more detailed mathematical analysis of density functions such as the one from Reid et al. [16] and those of equations (12) and (20). We note that in the interest of mathematical generality, in this appendix we adopt slightly different notation than is used in Reid et al. [16] and section 2.

We begin with the observation that the density function (3) applied to the configuration from Reid et al. [16] can be written as

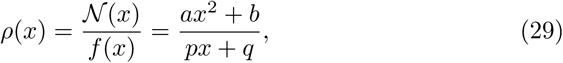

where *a, b, p, q* are parameters. The biological interpretation of equation (29) is that the quadratic function *𝒩*(*x*) = *ax*^2^ + *b* describes the relevant number of ants while the linear function *f* (*x*) = *px* + *q* describes the relevant linear distance of travel. The only *a priori* assumption that we place on the coefficients *a*, *b*, *p*, *q* is that *p*, *q* must be chosen so that *f* (*x*) = *px* + *q* is positive for all biologically reasonable values of the independent variable *x*. As discussed in Reid et al. [16], the fact that *𝒩*(*x*) is quadratic, while *f* (*x*) is linear and positive is a key point of the cost-benefit trade-off aspect of the theory of army ant living bridge formation.

We proceed with our analysis by computing the first and second derivatives of (29) with respect to the independent variable *x* thus obtaining

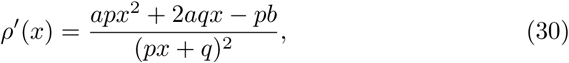

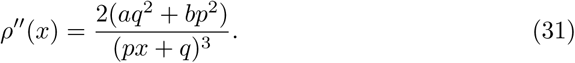

Now we seek to determine conditions under which equation (29) is maximized for a unique positive value *x*\* ∈ [0, *M*], where *M* represents the maximum possible distance value. Thus, we seek to determine a positive value of *x* in the interval [0, *M*] such that *ρ′*(*x*) = 0 and *ρ″*(*x*) < 0. Using the assumption that *f* (*x*) = *px* + *q* is positive for all biologically reasonable values of the independent variable *x*, this will happen whenever *apx*^2^ + 2*aqx* – *pb* = 0 and *aq*^2^ + *bp*^2^ < 0, and therefore whenever *x*\* satisfies

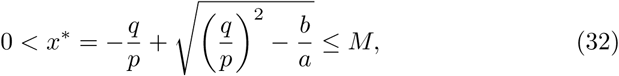

and

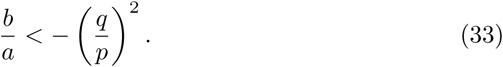

Note that in order to obtain a positive maximizing value of *x* in the interval [0, *M*], it must be the case that 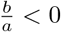.

Conditions (32) and (33) can easily be used to recover the results on optimal bridge positioning from Reid et al. [16]. The benefit of the different approach taken here is that it is applicable in situations not necessarily covered by the analysis of Reid et al. [16], provided that the configuration is such that the positioning of the army ant living bridge is completely determined by a single distance variable *x*. More interestingly, the analysis just given suggests how to move to multi-variable problems via analogy.

Consider the two-variable function

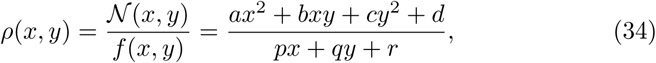

where now the only assumptions on the parameters *a*, *b*, *c*, *d*, *p*, *q*, *r* is that *p*, *q*, *r* are such that *f* (*x*) = *px* + *qy* + *r* is positive for all biologically reasonable values of the independent variables *x*,*y*. We note two points regarding equation (34): While we restrict our analysis to the two-variable case for notational simplicity, our work makes clear how to proceed in cases of three or more variables. More importantly, while (34) is similar in form to equations (12) and (20) of section 2; it is only locally equivalent due to the presence of the square root in (11) and the absolute value in (24). Nevertheless, an analysis of (34) still provides valuable insight into the results we obtain from equations (12) and (20), namely it aids in the explanation for the symmetry of the results derived from (20).

As before, we proceed with our analysis by computing the first and second derivatives of (34) with respect to the independent variables *x*, *y* thus obtaining

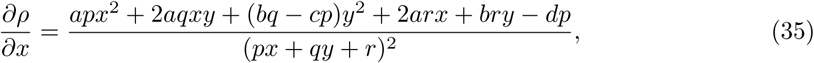

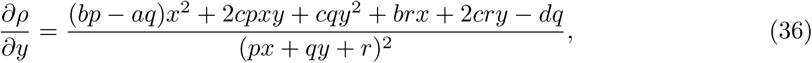

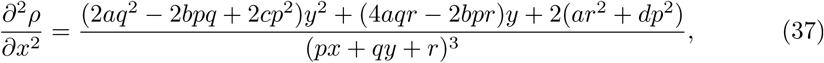

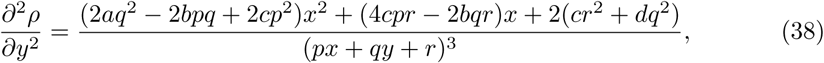

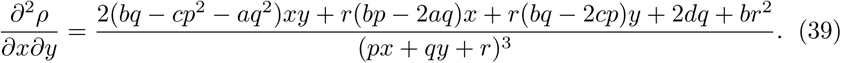

We would again like to find conditions for unique positive values for *x* and *y* in intervals [0, *M*] and [0, *N*] respectively that maximize the function (34)). In general, the complexity of the expressions in equations (35)-(39) make solving explicitly for maximizing values of *x* and *y* difficult. However, there are simplifying assumptions that can be made that are relevant to the multi-bridge configurations studied in section 2, that is, the case whenever *b* = 0, *a* = c, and *p* = q. This is in perfect analogy with the density function (20) from section 2. Under these assumptions, we get

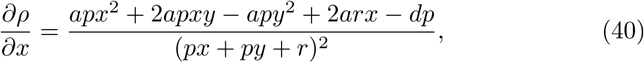

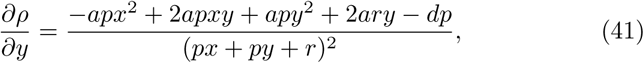

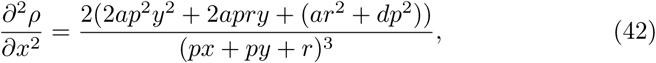

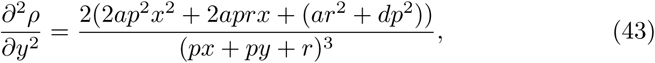

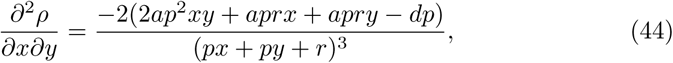

from which one can see that there is a value *t* satisfying *x* = *y* = *t* and

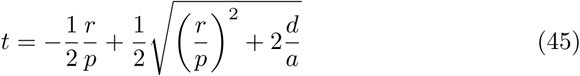

so that 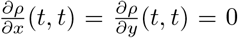. That is, there is a symmetric critical point for equation (34). Moreover, it is easy to see that when evaluated at (*t*,*t*) we have

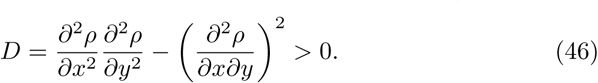

Thus, if 2*ap*^2^*t*^2^ + *2aprt* + *ar*^2^ + *dp*^2^ < 0, then the symmetric critical point (*t*,*t*) is at least a local maximum for (34). In addition, one can conclude from (45) when *t* will be in an interval of the form [0, *M*]. This analysis aids in our understanding of the symmetry of the results summarized in figure 6(b) obtained for the optimal bridge-position in the two-bridge configuration such as illustrated by figure 3. Similar reasoning for three independent variables can help to explain the symmetric results for the three-bridge configuration.

Now, we derive the results for the constrained optimization problem described in section 3. Specifically, we derive what is predicted to happen when the overall living bridge length is interpreted as a parameter. Doing so provides insight into the process of optimal bridge formation. Let *b_f_* represent the bridge length parameter. When *b_f_* is fixed, through equations (10)-(12) we arrive at a constrained optimization problem. That is, we seek to optimize the density

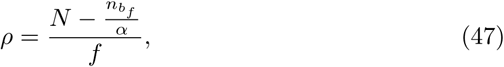

subject to the constraint

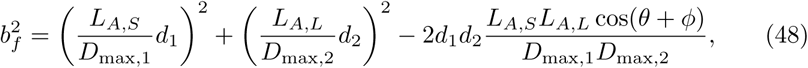

where

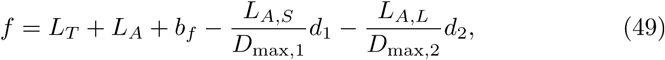

and

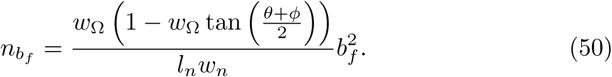

We note that *_n_f__* and hence 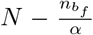 are now held constant. The values for *L_A,S_*, *L_A,L_, D*_max,1_ and *D*_max,2_ are obtained just as before.

Examining equations (47) and (48) we see that we need to maximize a function of the form

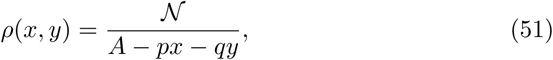

subject to a constraint of the form

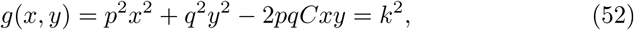

where *𝒩*, *A*, *k*, *p*, *q* and *C* are parameters. To simplify the problem, we observe that maximizing (51) subject to (52) is equivalent to minimizing *f* (*x*,*y*) = *A* — *px* — *qy* subject to the same constraint. This is done in a straightforward manner using the method of Lagrange multipliers. That is, we solve

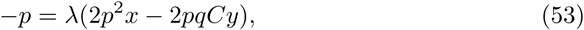

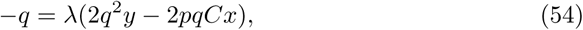

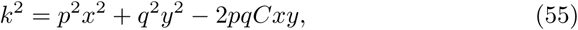

for λ, *x* and *y* that minimize *f* (*x*, *y*) = *A* – *px* — *qy.* This is easily done using (53) and (54) to set *px* – *qCy* = *qy – pCx* and then substituting into (55) and solving for the remaining variable. This gives solution

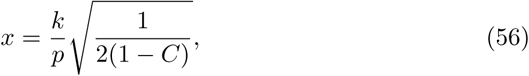

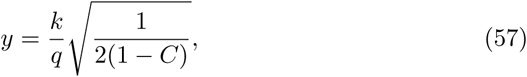

where we have retained only the positive square roots since in our application we seek positive distance values. Setting *k* = *b_f_*, 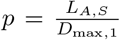, 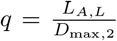 and *C* = cos(*θ* + *ϕ*)

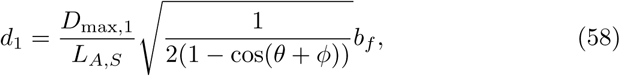

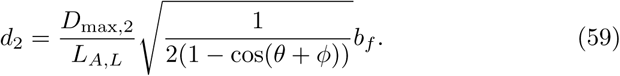

Keep in mind that we must set *d_i_* = *D*_max, *i*_ if *b_f_* is such that the predicted value of either *d*_1_ or *d*_2_ is greater than or equal to *D*_max,1_ or *D*_max,2_ respectively. Furthermore, using the expressions (8) and (9) together with equations (58) and (59), we see that the ratio of optimal distance values *d*_1_ and *d*_2_ satisfies

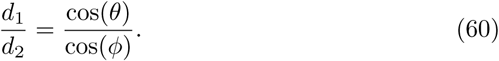

From this equation we can deduce interesting predictions. In particular, rearranging equation (60) gives

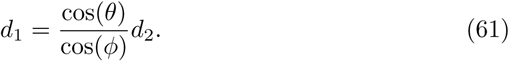

The biological consequences of this are described in 3.1.

## Acknowledgments

ABK is supported by a James S. McDonnell Foundation Postdoctoral Fellowship Award in Studying Complex Systems. DAW’s effort was supported through the Mathematical Biosciences Institute’s Research Experience for Undergraduates Program (NSF DMS - 1461163).

## Footnotes

2 Although a slight abuse of notation, throughout section 3 we use *d*_1_, etc. to denote not variables, but the value of the distance variables that actually maximize the relevant density function *ρ*.

## References

[1] S. Camazine, J.-L. Deneubourg, N. R. Franks, J. Sneyd, G. Theraulaz, E. Bonabeau, Self-Organization in Biological Systems, Princeton University Press, Princeton, NJ, ISBN 9780691116242, 2001.

[2] S. Garnier, J. Gautrais, G. Theraulaz, The biological principles of swarm intelligence, Swarm Intelligence 1 (1) (2007) 3–31, ISSN 1935-3812, 1935-3820, URL http://link.springer.com/article/10.1007/s11721-007-0004-y.

[3] I. D. Couzin, J. Krause, Self-Organization and Collective Behavior in Vertebrates, Advances in the Study of Behavior 32 (2003) 1–75, ISSN 0065-3454, URL http://www.sciencedirect.com/science/article/pii/S0065345403010015.

[4] I. D. Couzin, Collective cognition in animal groups, Trends in cognitive sciences 13 (1) (2009) 36–43, ISSN 1364–6613, URL http://www.ncbi.nlm.nih.gov/pubmed/19058992.

[5] D. J. T. Sumpter, Collective Animal Behavior, Princeton University Press, Princeton, NJ, ISBN 9780691129631, 2010.

[6] V. Guttal, I. D. Couzin, Social interactions, information use, and the evolution of collective migration, Proceedings of the National Academy of Sciences of the United States of America 107 (37) (2010) 16172–16177, ISSN 0027-8424, 1091-6490, URL http://www.pubmedcentral.nih.gov/articlerender.fcgi?artid=2941337&tool=pmcentrez&rendertype=abstract.

[7] M. Wolf, R. H. J. M. Kurvers, A. J. W. Ward, S. Krause, J. Krause, Accurate decisions in an uncertain world: collective cognition increases true positives while decreasing false positives, Proceedings. Biological sciences / The Royal Society 280 (1756) (2013) 20122777, ISSN 0962-8452, 1471-2954, URL http://dx.doi.org/10.1098/rspb.2012.2777.

[8] I. D. Couzin, J. Krause, N. R. Franks, S. A. Levin, Effective leadership and decision-making in animal groups on the move, Nature 433 (7025) (2005) 513–516, ISSN 0028-0836, 1476-4687, URL http://www.nature.com/nature/journal/v433/n7025/abs/nature03236.html.

[9] A. Duarte, F. J. Weissing, I. Pen, L. Keller, An Evolutionary Perspective on Self-Organized Division of Labor in Social Insects, Annual review of ecology, evolution, and systematics 42 (1) (2011) 91–110, ISSN 1543-592X, URL http://dx.doi.org/10.1146/annurev-ecolsys-102710-145017.

[10] C. Detrain, J.-L. Deneubourg, Self-organized structures in a superorganism: do ants “behave” like molecules?, Physics of life reviews 3 (3) (2006) 162–187, ISSN 1571-0645, URL http://www.sciencedirect.com/science/article/pii/S1571064506000200.

[11] G. Theraulaz, E. Bonabeau, S. C. Nicolis, R. V. Solé, V. Fourcassié, S. Blanco, R. Fournier, J.-L. Joly, P. Fernández, A. Grimal, P. Dalle, J.-L. Deneubourg, Spatial patterns in ant colonies, Proceedings of the National Academy of Sciences of the United States of America 99 (15) (2002) 9645–9649, ISSN 0027-8424, URL http://www.pubmedcentral.nih.gov/articlerender.fcgi?artid=124961&tool=pmcentrez&rendertype=abstract.

[12] J. Buhl, J.-L. Deneubourg, A. Grimal, G. Theraulaz, Self-organized digging activity in ant colonies, Behavioral ecology and sociobiology 58 (1) (2005) 9–17, ISSN 0340-5443, URL http://www.springerlink.com/index/10.1007/s00265-004-0906-2.

[13] H. King, S. Ocko, L. Mahadevan, Termite mounds harness diurnal temperature oscillations for ventilation, Proceedings of the National Academy of Sciences of the United States of America 112 (37) (2015) 11589–11593, ISSN 0027-8424, 1091-6490, URL http://dx.doi.org/10.1073/pnas.1423242112.

[14] A. Berdahl, C. J. Torney, C. C. Ioannou, J. J. Faria, I. D. Couzin, Emergent Sensing of Complex Environments by Mobile Animal Groups, Science 339 (6119) (2013) 574–576, ISSN 0036-8075, URL http://science.sciencemag.org/content/339/6119/574.

[15] P. C. Cross, E. K. Cole, A. P. Dobson, W. H. Edwards, K. L. Hamlin, G. Luikart, A. D. Middleton, B. M. Scurlock, P. J. White, Probable causes of increasing brucellosis in free-ranging elk of the Greater Yellowstone Ecosystem, Ecological Applications 20 (1) (2010) 278–288, ISSN 1939-5582, URL http://dx.doi.org/10.1890/08–2062.1.

[16] C. R. Reid, M. J. Lutz, S. Powell, A. B. Kao, I. D. Couzin, S. Garnier, Army ants dynamically adjust living bridges in response to a cost-benefit trade-off, Proceedings of the National Academy of Sciences 112 (49) (2015) 15113–15118.

[17] T. C. Schneirla, The army-ant behavior pattern: nomad-statary relations in the swarmers and the problem of migration, The Biological bulletin 88 (2) (1945) 166–193, ISSN 0006-3185, URL http://www.jstor.org/stable/10.2307/1538043.

[18] C. W. Rettenmeyer, Behavioral studies of army ants. Estudios de comportamiento de hormigas guerreras, The University of Kansas Science Bulletin. 44 (9) (1963) 281–465.

[19] T. C. Schneirla, Army Ants: A Study in Social Organization, W.H.Freeman & Co Ltd, 1972.

[20] J.-L. Deneubourg, S. Aron, S. Goss, J. M. Pasteels, The self-organizing exploratory pattern of the argentine ant, Journal of insect behavior 3 (2) (1990) 159–168, ISSN 0892-7553, URL http://www.springerlink.com/index/L76621W5417109QQ.pdf.

[21] C. Devigne, C. Detrain, Collective exploration and area marking in the ant Lasius niger, Insectes sociaux 49 (4) (2002) 357–362, ISSN 0020-1812, URL http://www.springerlink.com/index/qpern5ccjkld2d5e.pdf.

[22] C. Kost, E. G. de Oliveira, T. A. Knoch, R. Wirth, Spatio-temporal permanence and plasticity of foraging trails in young and mature leaf-cutting ant colonies (Atta spp), Journal of tropical ecology 21 (06) (2005) 677, ISSN 0266-4674, URL http://www.journals.cambridge.org/abstract_S0266467405002592.

[23] J. J. Howard, Costs of trail construction and maintenance in the leaf-cutting ant Atta colombica, Behavioral ecology and sociobiology 49 (5) (2001) 348–356, ISSN 0340-5443, URL http://link.springer.com/10.1007/s002650000314.

[24] A. I. Bruce, M. Burd, Allometric scaling of foraging rate with trail dimensions in leaf-cutting ants, Proceedings of the Royal Society B: Biological Sciences 279 (1737) (2012) 2442–2447, ISSN 1471-2954, URL http://www.ncbi.nlm.nih.gov/pubmed/22337696.

[25] T. Bochynek, B. Meyer, M. Burd, Energetics of trail clearing in the leaf-cutter ant Atta, Behavioral ecology and sociobiology 71 (1) (2016) 14, ISSN 0340-5443, 1432-0762, URL http://link.springer.com/article/10.1007/s00265-016-2237-5.

[26] R. V. Solé, E. Bonabeau, J. Delgado, P. Fernández, J. Marín, Pattern formation and optimization in army ant raids, Artificial life 6 (3) (2000) 219–226, ISSN 1064-5462, URL http://www.ncbi.nlm.nih.gov/pubmed/11224916.

[27] C. Anderson, G. Theraulaz, J.-L. Deneubourg, Self-assemblages in insect societies, Insectes sociaux 49 (2) (2002) 99–110, ISSN 0020-1812, URL http://www.springerlink.com/openurl.asp?genre=article&id= doi:10.1007/s00040-002-8286-y.

[28] S. Garnier, T. Murphy, M. Lutz, E. Hurme, S. Leblanc, I. D. Couzin, Stability and responsiveness in a self-organized living architecture, PLoS Comput Biol 9 (3) (2013) e1002984.

[29] S. Powell, N. R. Franks, How a few help all: living pothole plugs speed prey delivery in the army ant Eciton burchellii, Animal Behaviour 73 (6) (2007) 1067–1076.

[30] S. Powell, N. R. Franks, Caste evolution and ecology: a special worker for novel prey, Proceedings of the Royal Society of London B: Biological Sciences 272 (1577) (2005) 2173–2180, URL http://rspb.royalsocietypublishing.org/content/272/1577/2173.

[31] S. Powell, N. R. Franks, Ecology and the evolution of worker morphological diversity: a comparative analysis with Eciton army ants, Functional Ecology 20 (6) (2006) 1105–1114, ISSN 1365-2435, URL http://dx.doi.org/10.1111/j.1365-2435.2006.01184.x.

[32] D. Ardia, J. O. Arango, N. G. Gomez, Jump-Diffusion Calibration using Differential Evolution, Wilmott Magazine 55 (2011) 76–79, URL http://www.wilmott.com/.

[33] D. Ardia, K. Boudt, P. Carl, K. M. Mullen, B. G. Peterson, Differential Evolution with DEoptim: An Application to Non-Convex Portfolio Optimization, The R Journal 3 (1) (2011) 27–34, URL http://journal.r-project.org/archive/2011-1/2011-1_index.html.

[34] D. Ardia, K. M. Mullen, B. G. Peterson, J. Ulrich, DEoptim: Differential Evolution in R, URL http://CRAN.R-project.org/package=DEoptim, version 2.2-3, 2015.

[35] K. Mullen, D. Ardia, D. Gil, D. Windover, J. Cline, DEoptim: An R Package for Global Optimization by Differential Evolution, Journal of Statistical Software 40 (6) (2011) 1–26, URL http://www.jstatsoft.org/v40/i06/.

[36] K. V. Price, R. M. Storn, J. A. Lampinen, Differential Evolution - A Practical Approach to Global Optimization, Natural Computing, Springer-Verlag, iSBN 540209506, 2006.

[37] R Core Team, R: A Language and Environment for Statistical Computing, R Foundation for Statistical Computing, Vienna, Austria, URL https://www.R-project.org/, 2016.

